# A simple description of biological transmembrane potential from simple biophysical considerations: no equivalent circuits

**DOI:** 10.1101/2020.05.02.073635

**Authors:** Marco Arieli Herrera-Valdez

**Affiliations:** Systems Physiology Laboratory, Department of Mathematics, Facultad de Ciencias, Universidad Nacional Autónoma de México, CDMX, México

## Abstract

Biological membranes mediate different physiological processes necessary for life. Ion movement around, into and out of cells, is arguably one of the most important of such processes, as it underlies electrical signalling and communication within and between cells. The difference between the electrical potentials inside and outside a biological membrane, called transmembrane potential, or membrane potential for short, is one of the main biophysical variables affecting ionic movement. Most of the equations that describe the change in membrane potential are based on analogies with resistive-capacitative electrical circuits, with conductance and capacitance proposed in classical studies as measures of the permeable and the impermeable properties of the membrane, respectively. However, the parts in ohmic circuits are not like the biological elements present in the spaces around and within biological membranes. This article presents a simple derivation for an equation describing local changes in transmembrane potential that is not based on electrical circuit analogies. In doing so, concepts like the so-called “membrane capacitance” are explained and generalized. Importantly, the classical model for the membrane potential based on an equivalent RC-circuit is recovered as particular case from the general derivation presented here. Modeling examples are presented to illustrate the use of the derivation and the effects of changing the way charges aggregate around the membrane as a function of the membrane potential.

## 1 Introduction

Biological membranes mediate communication between cellular compartments and their surrounding environments (Blaustein et al., 2004; Boron and Boulpaep, 2016; Helman and Thompson, 1982; Sten-Knudsen, 2002). One of the most important biophysical variables affecting, and affected by such communication at the level of the plasma membrane, is the difference between the electrical potential inside and outside the membrane, or membrane potential (Johnston et al., 1995; Sperelakis, 2012; Weiss, 1996). Paraphrasing Cole (1933), and others (Fricke, 1931; Höber, 1936), the existence of the membrane potential is, by itself, an indication of the fact that a cell is alive, and changes in the membrane potential are physiological correlates of cellular activity. The membrane potential is thus a key determinant of electrical and biochemical signaling in different levels of biological organization, so basic understanding the biophysical principles underlying its behavior is important.

The transmembrane potential is generated by the presence of ions of different species across membranes (Briggs, 1930; Brooks, 1929), different permeabilities across the membrane (Cole, 1940; Höber, 1936), and the continuous passive and active transport contributing to electrical signaling and other physiological processes (Eisenberg, 1998, 1990; Hille, 1992; Höber et al., 1939; Hodgkin and Katz, 1949a,b). Ion transport around and across membranes is necessary for a variety of physiological processes including, but not limited to electrical signaling, secretion, and volume regulation. Importantly, also has an effect on the transmembrane potential.

Modeling approaches have been useful to describe electrical and biochemical interactions involving ions, and how they affect the membrane potential (Fricke, 1931; Osterhout, 1933; Teorell, 1935). Many models of biological membranes are based on constructing equivalent electrical circuits (Hermann, 1905, 1899) using conservative paradigms like Kirchoff’s law (Johnson et al., 1989). Of seminal importance, the Hodgkin and Huxley (1952) model describes the changes in membrane potential in a system of differential equations derived from an RC-circuit with resistors, a capacitor, and a battery arranged in parallel, representing membrane channels, its surrounding compartments, and the membrane potential. The total current in the model is the sum of a “‘capacitative” current representing electrical flow around the membrane, and different resistive currents representing different transmembrane ion fluxes mediated by ion channels (Hodgkin et al., 1952). Charged molecules do accumulate next to the hydrophilic heads of the lipid bilayer and the charge density around the membrane increases as the membrane potential increases. However, ionic transmembrane currents are not resistive (ions crossing the membrane are not like electrons through a wire) and the lipid bilayer of the membrane is not quite like the space between the plates in a capacitor. Also, in consideration of different proteins, structures, and functional environments around cells, the charge density around the membrane is not necessarily proportional to the membrane potential. To exacerbate the matter, there are multiple problems to experimentally measuring the membrane capacitance in cells (Golowasch et al., 2009).

It is possible to derive a description for the change in the transmembrane potential in a small patch of membrane by taking into account the charge densities of ions in and around the membrane, without using electrical circuit analogies. The result is a simple differential equation that explains the phenomenological formulation originally proposed by Hodgkin and Huxley (1952) as a particular case. Briefly, a general expression for the voltage-dependent change in the charge around the membrane can be simplified to obtain the so-called “membrane capacitance” by assuming that the charge density around the membrane is linearly related to the membrane potential. Interestingly, the term emerges from the chain-rule while considering the time-dependent change in the total charge density around the membrane and its dependence on voltage (Coombs et al., 1959; Everitt and Haydon, 1968; Lu et al., 1995). This is in line with the thinking behind the thermodynamic model for transmembrane transport by Herrera-Valdez (2018), a general formulation that describes different physiological transport mechanisms, both passive and active, that also includes conductance based models as particular cases.

Once the derivation is explained, it is illustrated with two simple examples of membrane excitability models.

## 2 Biophysical derivation to describe the transmembrane potential

Consider a small volume Ω containing a small portion of a cellular membrane, together with the extracellular and intracellular compartments in the immediate neighborhood around the membrane (Fig. 1). Assume, each of these compartments is filled with physiological solution containing ions of different species, and assume also that the number of ions for the different ion species is constant. By extension, the total number of ions in the system and the total charge (in Coulombs) are constant in time too. Assume also that each of the ion types has some permeability across the membrane and as a consequence, there is a time-dependent charge density *Q*_*m*_(*t*) in the space within the lipid bilayer of the membrane.

**Figure 1:**
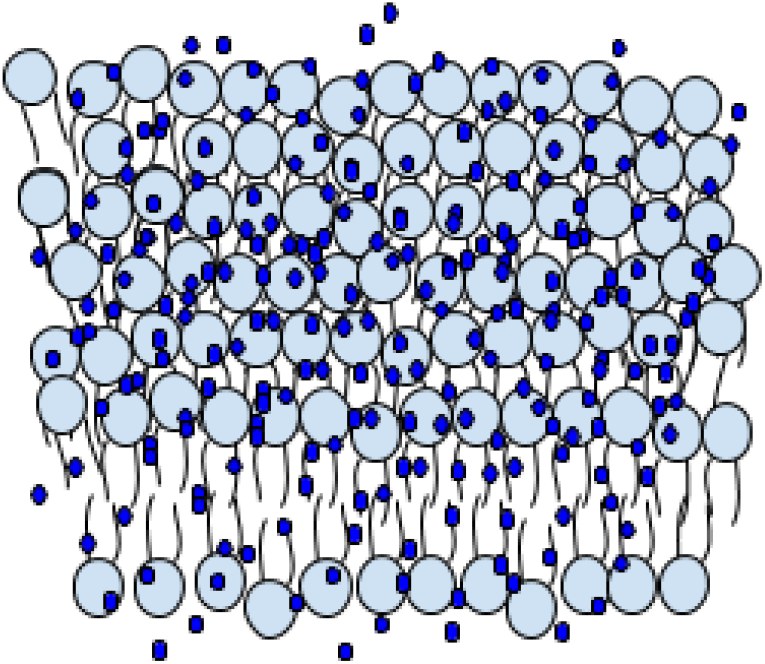
Plasma membrane with ions around and within the space in the lipid bilayer of the membrane.

Let *U*_*e*_(*t*) and *U*_*i*_(*t*) (in Volts) represent the electrical potential in the extra- and intracellular compartments, respectively. The difference *v*(*t*) = *U*_*i*_(*t*) − *U*_*e*_(*t*) is the transmembrane potential, or by simplicity, membrane potential. For conceptual simplification, if the extracellular potential is equal to zero, then the transmembrane potential can be thought of as the intracellular potential. Since ions tend to accumulate around the membrane as *v* increases, the net charge density (Coulombs/*μ*m^2^) around the membrane can be assumed to be some smooth, monotonic, increasing function *Q*_*a*_(*v*) of the membrane potential that changes its sign if the value of *v* changes from negative to positive. As a consequence, time variations in *v* would then induce time variations in *Q*_*a*_.

It follows from the assumption that the total charge *Q*_*a*_ + *Q*_*m*_ is constant, that the time-dependent change in the total charge density is zero. That is,

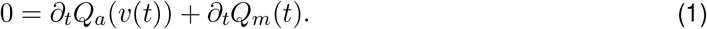

Here *∂*_*t*_ represents the instantaneous rate of change with respect to time. Therefore, term *∂*_*t*_*Q*_*m*_(*t*) represents the current density (Amperes/*μ*m^2^) carried by different ions across the membrane and *∂*_*t*_*Q*_*a*_(*v*(*t*)) represents the current density around the membrane.

Equation (1) yields a differential equation for *v* in terms of the ionic flux across the membrane, normalized by the change in charge density around the membrane as a function of *v*. To see this, note first that applying the chain rule to the term representing the charge density around the membrane yields

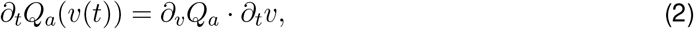

with *∂*_*t*_*v* in V/s and *∂*_*v*_*Q*_*a*_ in Coulombs/*μ*m^2^ per Volt. Also, the current density across the membrane can be thought of as proportional to the total transmembrane flux of charge. Separating terms in equation (1) and solving for *∂*_*t*_*v* from equation (2) yields

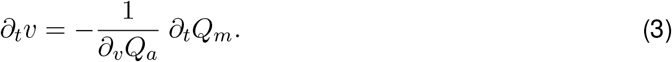

where *∂*_*t*_*Q*_*m*_(*t*) represents the total transmembrane current density carried by the different ion types around the membrane (Amperes/*μ*m^2^).

To build specific models, the transmembrane flux *∂*_*t*_*Q*_*m*_ can be substituted by a sum of different transmembrane ionic fluxes associated to different physiological transport mechanisms (Herrera-Valdez, 2018).

The inverse of *∂*_*v*_*Q*_*a*_ can be thought of as a scaling factor for the change in the membrane potential. If the profile for *Q*_*a*_(*v*) is nonlinear, then the scaling varies dynamically, possibly amplifying or dampening the way that *v* changes in time. One way to explore this possibility is to examine how the profile of charge accumulation around the membrane affects the dynamics of a neuronal action potential. To do so, let us define three possible different profiles for the charge density around the membrane (*Q*_*a*_) as a function of *v*, and note an important particular case that gives theoretical validation to the equivalent circuit analogy for membranes.

### 2.1 Linear accumulation of charge around the membrane and the membrane capacitance from the equivalent circuit models

One particular case of importance can be formulated by assuming that charge aggregation around the membrane is linear as a function of voltage. That is, *Q*_*a*_(*v*) = *C*_*m*_*v*, where *C*_*m*_ (in pF/*μ*m^2^) is a constant that describes the voltage-dependent rate of charge density around the membrane (Fig. 2A, black line). In this case, *∂*_*v*_*Q*_*a*_ = *C*_*m*_, which means that equation (3) is then 

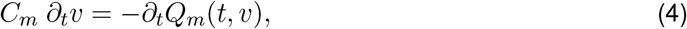

as proposed by Hermann, Fricke, Cole, and Hodgkin and Huxley, among others (Fig. 2B, black line).

**Figure 2:**
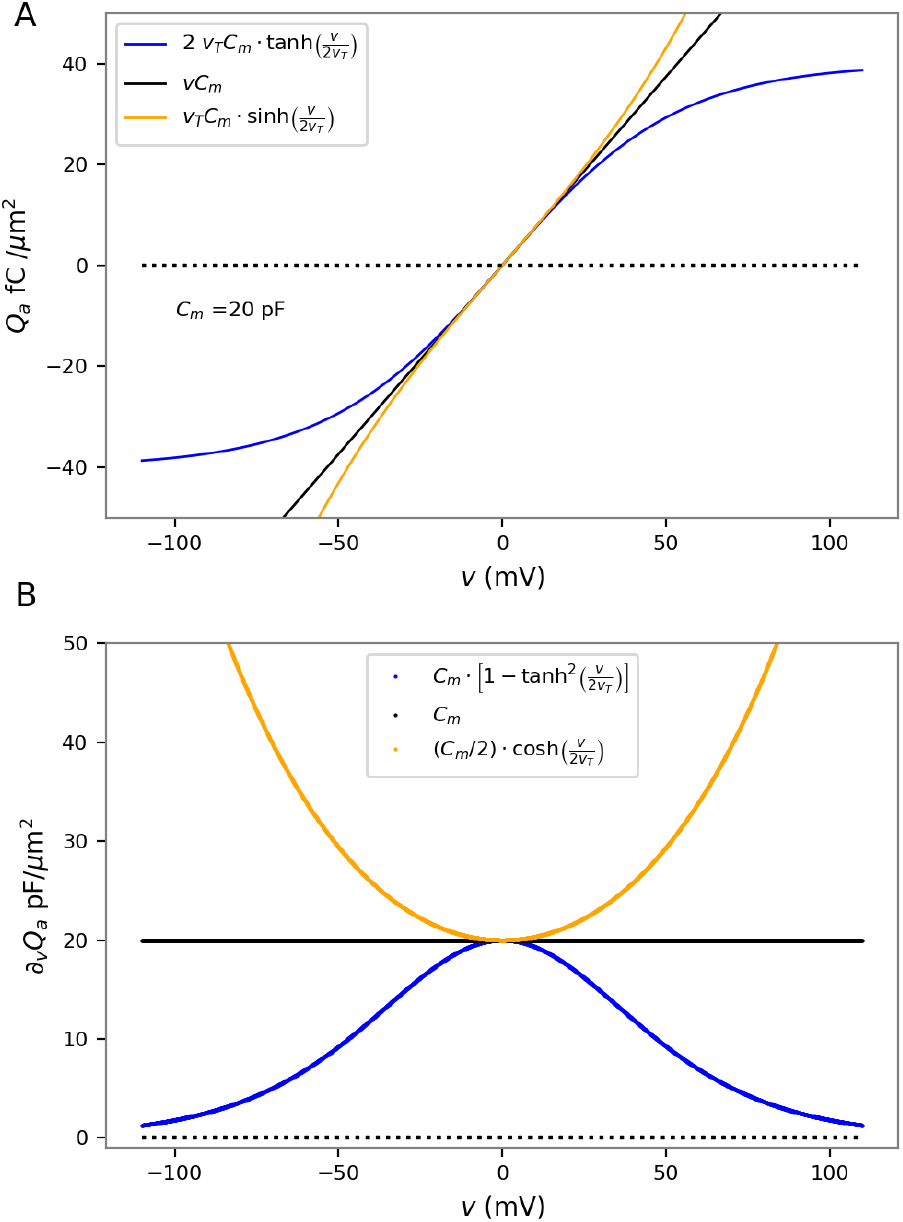
(A) Profiles for charge densities around the membrane and (B) their changes as a function of voltage (bottom).

The change in charge density around the membrane, *∂*_*v*_*Q*_*a*_, is in Coulombs/Volt per *μ*m^2^, equivalent to Farads/*μ*m^2^, the same units for capacitance. The plates of a capacitor in an electrical circuit are made of conducting media (e.g. metal), and separated by air. Therefore, this particular case supports the idea that the intra and extracellular media can be thought of as similar to the metal plates, and the membrane can be regarded as analogue to the insulating air layer between the plates.

### 2.2 Two nonlinear voltage dependent profiles for the density of charge around the membrane

#### *Q*_*a*_ with saturation as a function of *v*

It is arguable that there is an upper bound for the accumulation of charges around the membrane (Fig. 2A, blue curve) as a function of voltage. That is, if the membrane is polarized to very large voltages (either negative or positive) the charge density around the membrane should reach a limit. If this is the case, then the dependence of *Q*_*a*_ on voltage would be given by

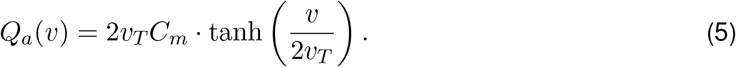

which means that,

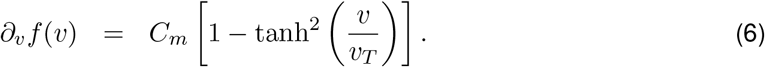

The graph of *Q*_*a*_(*v*) is odd and increasing, bounded by the asymptotic values of ±2*C*_*m*_, respectively (Fig. 2A, blue curve). The graph of *∂*_*v*_*Q*_*a*_ in equation (6) has a symmetrical shape around a local maximum at *v* = 0, always taking positive values (Fig. 2B, blue curve). This means that the charge density around the membrane does not change much when the membrane is polarised, and its maximum change occurs when the membrane is not polarised. As a consequence, *∂*_*v*_*Q*_*a*_ exerts an amplification effect on *∂*_*t*_*v* that increases as *v* is further from zero, especially near the resting potential of the cell.

#### Exponentially increasing charge density around the membrane

Another possibility similar to the current densities from the thermodynamic model (Herrera-Valdez, 2018), is that *Q*_*a*_(*v*) is a hyperbolic sine. In this case,

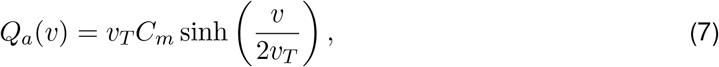

and

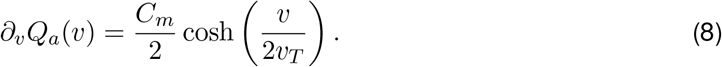

The graph of *Q*_*a*_(*v*) is odd and increasing with exponential growth in both directions of the *v* axis (Fig. 2A, orange curve), qualitatively opposite to that described earlier for the saturating *Q*_*a*_ with hyperbolic tangent shape. In this case *∂*_*v*_*Q*_*a*_ has a “U” shape with a local minimum of *C*_*m*_ at *v* = 0, which means that *∂*_*v*_*Q*_*a*_ exerts an attenuation effect on *∂*_*t*_*v* that increases as *v* is further from 0 (Fig. 2B, orange curve).

The nonlinear dependence of *∂*_*v*_*Q*_*a*_ from equations (6) and (8) could produce very different effects on the dynamics of the transmembrane potential.

Note the three profiles for the charge density around the membrane presented above can be thought of as approximations of one another around *v* = 0.

### 2.3 Neuronal dynamics assuming voltage dependent density of charge around the membrane

Consider a system with three transmembrane transport mechanisms, say, voltage-dependent K^+^ and Na^+^ channels, and a Na^+^-K^+^ ATPase. If *w* represents the proportion of open K^+^ channels and the proportion of inactivated Na^+^ channels, then the dynamics of the membrane potential can then be written as

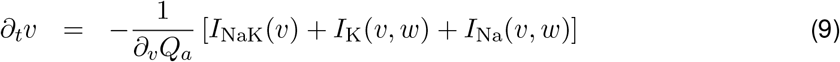

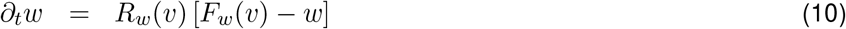

with transmembrane currents (pA) given by

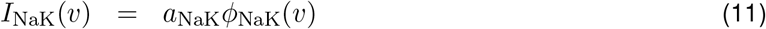

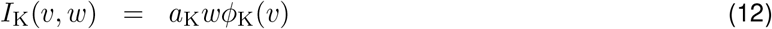

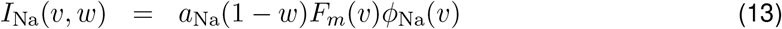

and auxiliary functions for the transmembrane fluxes, the activation rates and the steady state activation of the K^+^ channels given by

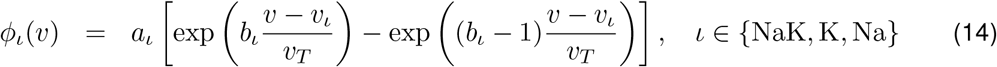

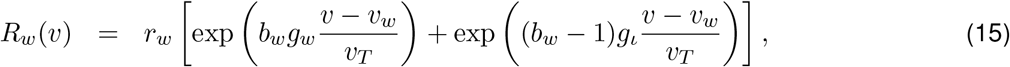

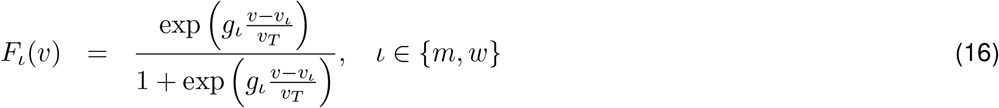

A description of all the parameters and their values can be found in Table 1.

**Table 1:**
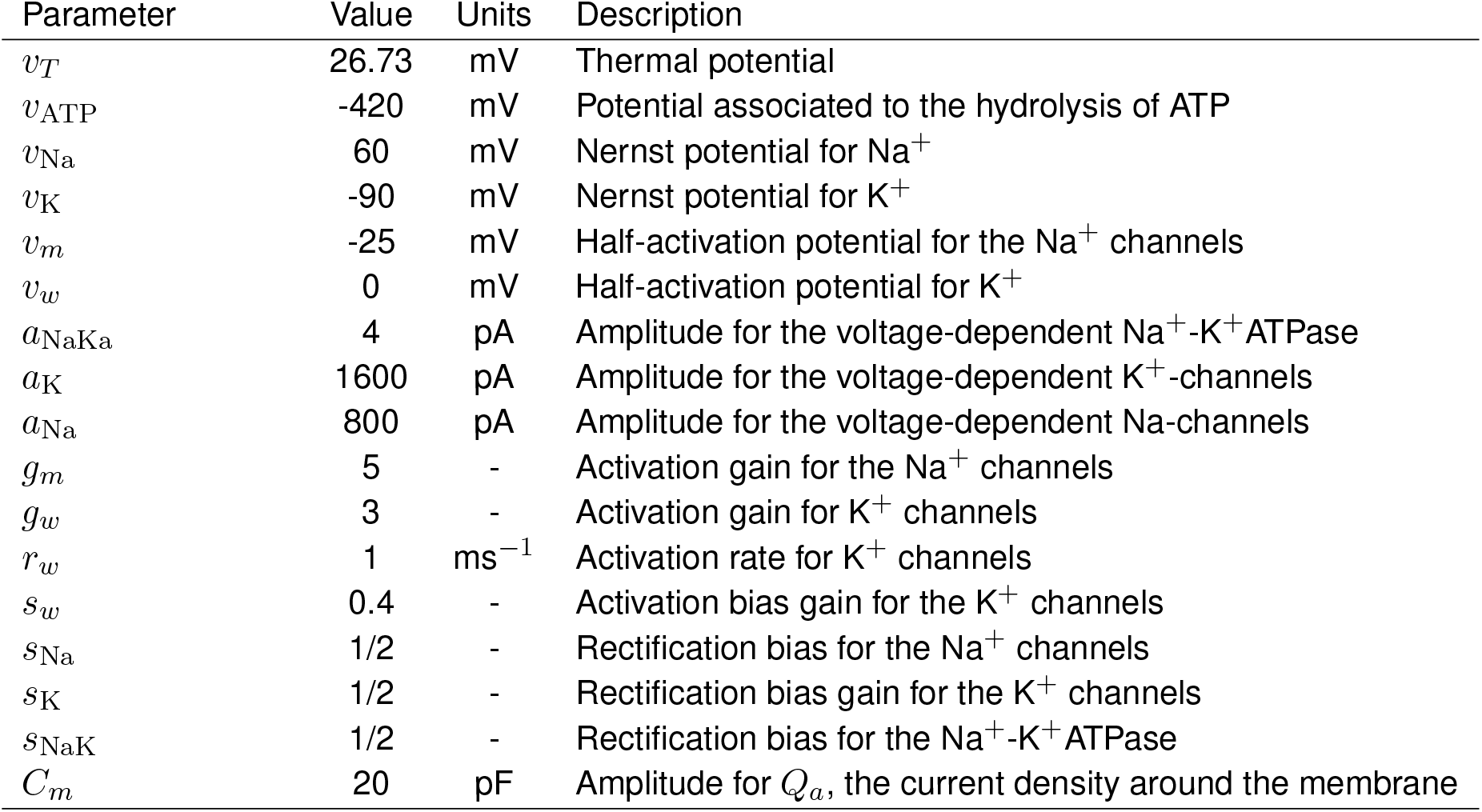
Parameters for the neuronal dynamics model. The reversal potential for the Na^+^-K^+^ATPase is given by *v*_NaK_ = *v*_ATP_ + 3*v*_Na_ - 2*v*_K_. Initial conditions at *v*_0_ = −48 mV and *w*_0_=0.001.

The dynamics for the transmembrane potential in a neuron were modelled for to compare the effects caused by the three profiles (tanh, linear, and sinh) for t he d ensity o f c harge a round the membrane *Q*_*a*_ (Fig. 3, blue, black, and orange lines, respectively).

**Figure 3:**
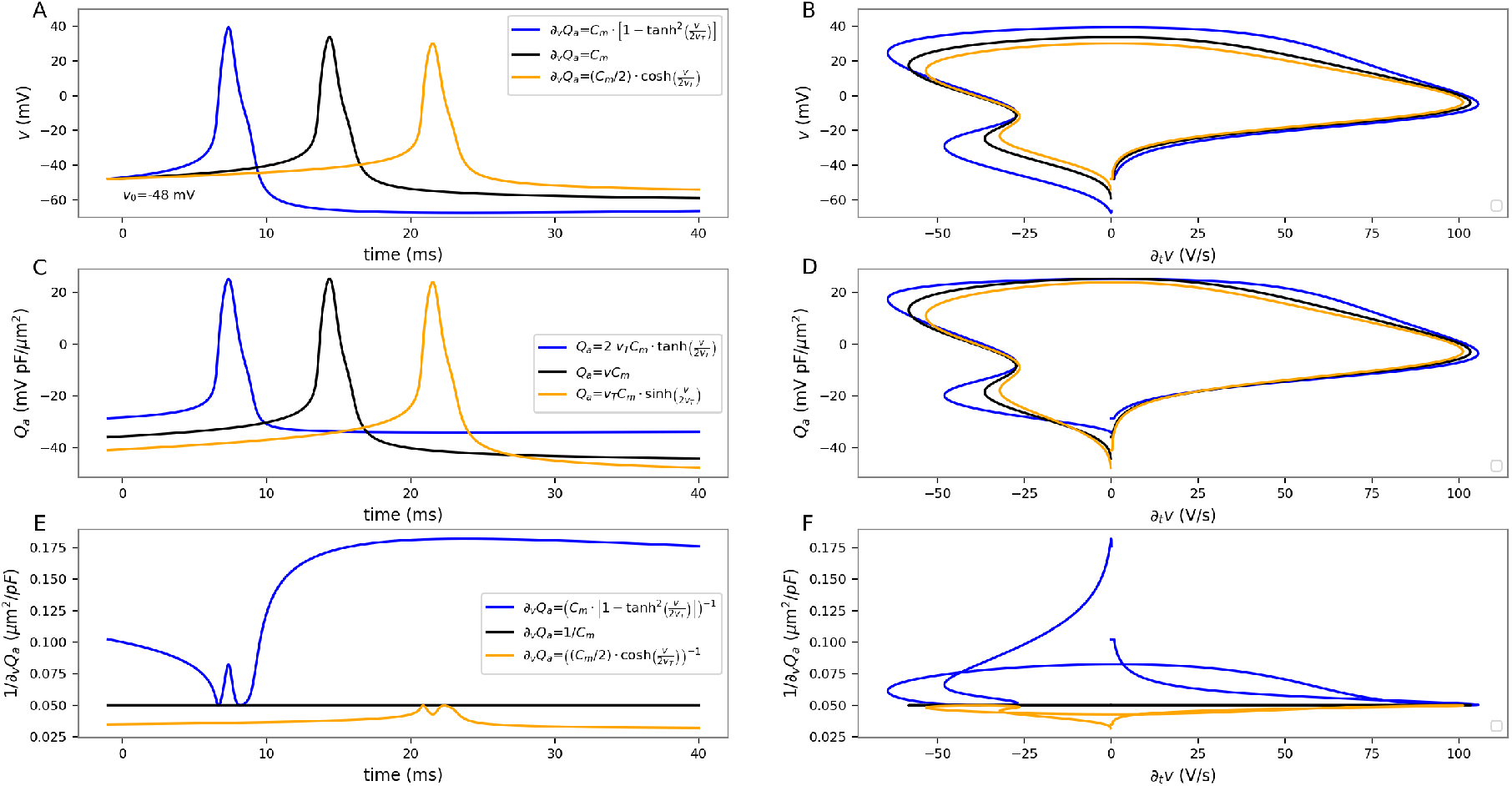
Effects of voltage-dependent scaling by the change in charge density around the membrane, on the dynamics of a neuronal action potential. The panels on the left column (A,C,E) show time series for *v*, *Q*_*a*_, and 1*/∂*_*v*_*Q*_*a*_, respectively. The panels on the right columns (B,D,F) show the corresponding trajectories of the same variables as a function of the voltage-dependent scaling induced by 1*/∂*_*t*_*v* on the instantaneous change in *v* with respect to time.

Note that the ranges for the three different profiles for *∂*_*v*_*Q*_*a*_ (Fig. 2) can be ordered by magnitude. The hyperbolic profile for *Q*_*a*_ yields the lowest values for *∂*_*v*_*Q*_*a*_, followed by the constant *∂*_*v*_*Q*_*a*_ = *C*_*m*_, surpassed by the hyperbolic sine profile that yields the largest values for *∂*_*v*_*Q*_*a*_. Therefore, the scaling effects of 1/*∂*_*t*_*Q*_*a*_ on the change in transmembrane potential can be thought of in the opposite order, with larger effects caused by *Q*_*a*_ with linear and saturating profiles, a smaller scaling for the linear profile, and the smallest scaling caused by the exponential profile. It is possible to appreciate this by comparing the trajectories of three neuronal action potentials from the same initial conditions (*v*_0_ = −48 mV and *w*_0_=0.001), obtained from a model with identical parameters (Herrera-Valdez et al., 2013) that only differ in their *Q*_*a*_ profile (Fig. 3).

The smallest scaling effect on the change in *v* occurs for the hyperbolic sine profile (Fig. 3, orange curves, spike time at 6.62 milliseconds approx), as shown by the longer delay in the action potential in comparison with the linear and tanh profiles (Fig. 3A-B, black and blue curves, respectively, spike times at 13.63 and 20.82 milliseconds approx). The delay can be increased if the initial condition is lower, or decreased if higher, without altering the hierarchy in the scaling effect. The smaller scaling effects caused by the sinh profile for *Q*_*a*_ on *∂*_*t*_*v*, in comparison to the linear and tanh-profiles for *Q*_*a*_, can also be appreciated in the trajectories of the (*∂*_*t*_*v, v*)-plane, which shows a smaller area contained by the curve for the sinh-profile, in comparison to the trajectories produced by the linear and tanh-profiles for *Q*_*a*_. Similar ranges for *Q*_*a*_ can be observed (Fig. 3C-D), but the dynamics show smaller departures from 0 in the tanh-profile with respect to the other two. The scaling induced by the inverse of *∂*_*v*_*Q*_*a*_ is notorious in both the time-dependence of the scaling factor, and also in its trajectory as a function of *∂*_*t*_*v* (Fig. 3E-F). The larger scaling effect of the tanh-profile on the time-dependent change in *v* can be clearly appreciated in Fig. 3F, which shows the values of the scaling factors of the tanh-profile during the action potential above any of the values from the linear or sinh profiles. To further illustrate the scaling effects of the *Q*_*a*_ profiles on *∂*_*t*_*v*, a small sample varying the initial conditions was calculated and the maximum upstroke velocities obtained in each case. In each case, it was possible to observe the larger amplification effect of the tanh and linear profiles (in that order) with respect to the sinh profile; in agreement with the observations made above.

One indirect way of measuring the effects of the scaling factor 1*/∂*_*t*_*Q*_*a*_ on the change in membrane potential, is to calculate the efficiency of the Na^+^current during the upstroke of the action potential. In brief, this is done by calculating the total Na^+^charge during the upstroke, and dividing it by the total charge during the same time interval. For the simulations shown above, the efficiencies increased for the tanh, linear, and sinh models respectively, reflecting the decreased effect of the scaling by 1*/∂*_*v*_*Q*_*a*_ on *∂*_*t*_*v* for the sinh profile (Table 2).

**Table 2:** Maximum rate of change for *v* with respect to time and efficiency of the Na^+^current during the upstroke of the action potential for the different profiles taking into account different starting voltages.

| $Q_a$ profile | $\tanh$ | $linear$ | $\sinh$ | $\tanh$ | $linear$ | $\sinh$ |
| --- | --- | --- | --- | --- | --- | --- |
| | $\max \partial_t v$ (V/s) | | | Na-efficiency | | |
| $v_0 = -46$ mV | 105.951 | 104.341 | 103.227 | 1.07043 | 1.10979 | 1.14603 |
| $v_0 = -47$ mV | 105.829 | 103.954 | 102.423 | 1.08322 | 1.13617 | 1.2005 |
| $v_0 = -48$ mV | 105.704 | 103.442 | 101.25 | 1.09777 | 1.18076 | 1.29603 |

## 3 Discussion

A simple derivation for an equation describing the time evolution of the transmembrane potential has been presented (equation (3)). The derivation is *not* based on an equivalent circuit analogy, but it is possible to obtain the traditional equation for the equivalent RC-circuit and explain the so-called "membrane capacitance" as a particular case; a desirable property of generalized models. The derivation draws from current knowledge about the structure of the membrane and its transmembrane proteins, and stems from a separating the ion-impermeable and ion-permeable aspects of the membrane, as referred by Cole and Curtis and other authors. The new equation is in line with the derivation of the the thermodynamic model (Herrera-Valdez, 2018), in that it only considers basic biophysical principles to describe the elements in a system that include a membrane, and ions in solution.

From a physical perspective, changes in the sign and magnitude of the transmembrane potential should cause changes in the density of charges around the membrane (Huang, 1981; Santos-Sacchi and Navarrete, 2002). Nevertheless, The membrane capacitance is typically regarded as a constant measure (Cole and Curtis, 1938). One important result from the derivation is that the change in the charge density around the membrane is not necessarily a constant, as initially found in early works (Everitt and Haydon, 1968), and evidenced sporadically over time in different reports (Amzica and Neckelmann, 1999; Neher and Marty, 1982; Proks and Ashcroft, 1995; Smith et al., 1997).

